# Diabetes induced decreases in PKA signaling in cardiomyocytes: the role of insulin

**DOI:** 10.1101/2020.04.02.021667

**Authors:** Craig A. Eyster, Satoshi Matsuzaki, Jennifer R. Giorgione, Kenneth M. Humphries

## Abstract

The cAMP-dependent protein kinase (PKA) signaling pathway is the primary means by which the heart regulates moment-to-moment changes in contractility and metabolism. We have previously found that PKA signaling is dysfunctional in the diabetic heart, yet the underlying mechanisms are not fully understood. The objective of this study was to determine if decreased insulin signaling contributes to a dysfunctional PKA response. To do so, we isolated adult cardiomyocytes (ACMs) from wild type and Akita type 1 diabetic mice. ACMs were cultured in the presence or absence of insulin and PKA signaling was visualized by immunofluorescence microscopy using an antibody that recognizes proteins specifically phosphorylated by PKA. We found significant decreases in proteins phosphorylated by PKA in wild type ACMs cultured in the absence of insulin. Akita ACMs also had decreased PKA signaling in the absence of insulin and this was not rescued by insulin. The decrease in PKA signaling was observed regardless of whether the kinase was stimulated with a beta-agonist, a cell-permeable cAMP analog, or with phosphodiesterase inhibitors. PKA content was unaffected, suggesting that the decrease in PKA signaling may be occurring by the loss of specific PKA substrates. Phospho-specific antibodies were therefore used to discern which potential substrates may be sensitive to the loss of insulin. Contractile proteins were phosphorylated similarly in wild type and Akita ACMs regardless of insulin. However, phosphorylation of the glycolytic regulator, PFK-2, was significantly decreased in an insulin-dependent manner in wild type ACMs and in an insulin-independent manner in Akita ACMs. These results demonstrate a defect in PKA activation in the diabetic heart, mediated in part by deficient insulin signaling, that results in an abnormal activation of a primary metabolic regulator.

## Introduction

Heart disease is the leading cause of death for patients with type I or type II diabetes [1]. This is in part because diabetes directly impacts cardiac function independently of other comorbidities. This is termed diabetic cardiomyopathy and it is a multi-factorial condition resulting from the metabolic stresses of disrupted insulin signaling, hyperglycemia and hyperlipidemia, and mitochondrial dysfunction [2]. In addition, there are also disruptions in protein kinase A (PKA) signaling, the molecular pathway that mediates the metabolic and contractile responses to sympathetic stimulation [3, 4]. While the molecular mechanisms contributing to diabetic cardiomyopathy are highly interrelated, the relationship between metabolic perturbances and changes in PKA signaling are not fully understood.

In the healthy heart the sympathetic nervous system functions through β-adrenergic signaling to increase cardiac contractility. Catecholamines bind to Gα_s_-coupled β-adrenergic receptors, stimulate adenylate cyclase, and subsequently increase cAMP to activate PKA. PKA then phosphorylates proteins involved in calcium cycling (troponin, SERCA, and phospholamban) and proteins that affect metabolic substrate selection (phosphofructokinase-2 (PFK-2) and acetyl-CoA carboxylase-2) [5, 6]. Glucose uptake and oxidation are the primary means of meeting the rapid increase in energy demands in response to sympathetic stimulation [6, 7]. In this way the increase in contractility is orchestrated with activation of metabolic pathways to ensure energy demands are met.

As an insulin sensitive tissue, the heart is affected by either decreases in circulating insulin or by the loss of insulin signaling that occur with type 1 or type 2 diabetes [8, 9]. The primary role of insulin is to increase glucose uptake and metabolism. Thus, the decrease in insulin signaling contributes to the metabolic inflexibility whereby the heart increases reliance on fatty acid oxidation, at the expense of decreased glucose usage, to meet energetic demands [10]. Over the long term, this metabolic inflexibility promotes lipotoxicity, mitochondrial dysfunction, and oxidative stress. Increasing evidence suggests there are interactions between insulin and β-adrenergic signaling. For example, hyperinsulinemia can blunt PKA signaling via an increase in phosphodiesterase 4 which increases cAMP hydrolysis [11, 12]. In our own work, we found PKA signaling is affected in a type 1 diabetic mouse model via changes in PKA activity that are downstream of receptor activation and adenylate cyclase activity [3]. Furthermore, we identified that a loss of insulin signaling, in both type 1 and type 2 diabetic conditions, decreases the content of the PKA substrate, PFK-2 [4]. In the healthy heart, phosphorylation of PFK-2 increases the production of fructose-2,6-bisphosphate, an allosteric activator of PFK-1 which is a committed and rate-limiting step of glycolysis [6]. Thus, the loss of insulin signaling disrupts a mechanism whereby β-adrenergic signaling increases glycolysis to meet energetic demands.

The goal of the present work was to define how the loss of insulin signaling impacts β-adrenergic signaling in cardiomyocytes. Adult mouse cardiomyocytes (ACMs) were isolated from control and Akita diabetic mice and then cultured in the presence or absence of insulin. ACMs were subsequently stimulated with β-adrenergic agonists and PKA signaling was determined by immunofluorescence microscopy. We have identified a striking decrease in PKA signaling in wild type ACMs cultured in the absence of insulin. This effect was mirrored in ACMs isolated from Akita type 1 diabetic mice, regardless of the presence of added insulin. Using phospho-specific antibodies, we found that the phosphorylation of proteins involved in calcium regulation were unaffected by the absence of insulin. In contrast, the metabolic target, PFK-2, was highly sensitive. Our results demonstrate the effects of PKA on cardiomyocyte function is dependent upon the actions of insulin.

## Materials and Methods

### Adult Mouse Cardiomyocyte Isolation

Adult cardiomyocytes from C57BL/6J or C57BL/6J-*Ins2^Akita^*/J mice (Akita, The Jackson Laboratory 003548) were isolated and cultured as previously described [3, 13]. Briefly, after isoflurane administration the heart was excised, the aorta was cannulated, and it was then perfused with type II collagenase (Worthington #LS004176). Calcium was reintroduced to the subsequent single cell suspension and cells were plated on laminin (Corning 354232) coated plates. Media was switched to serum-free culture media (minimal essential medium with Hanks’ balanced salt solution, Gibco (11575-032) supplemented with 0.2mg/mL sodium bicarbonate, penicillin-G, 0.1%BSA, glutamine, and 10mM butanedione monoxime. Cells were cultured 18h at 37°C and 5%CO_2_ with indicated drugs as described in figure legends. All procedures were approved by the Oklahoma Medical Research Foundation Animal Care and Use Committee.

### Antibodies and Drugs

Rabbit polyclonal antibodies to phospho-PKA substrate (9621S), PKA C-α (4782S), phospho-PFK2 (13064S), phospho-Ser16/Thr17-phospholamban (8496S), and phospho-Troponin I were purchased from Cell Signaling Technology. Rabbit polyclonal anti-PDE4D (ab14613) was purchased from Abcam. Alexa Fluor 488 goat anti-rabbit IgG (A11034) and Alexa Fluor 546 phalloidin (A22283) were purchase from Invitrogen. Insulin solution human (19278), (-)-Isoproterenol hydrochloride (16504), 3-Isobutyl-1-methylxanthine (15879) and 8-Bromoadenosine 3’,5’-cyclic monophosphate sodium salt (B7880) were purchased from Sigma. Phosphodiesterase inhibitor Tocriset containing Milrinone, Cilostamide, Zardaverine, *(R)-(-)-* Rolipram, and Ro 20-1724 (Cat. No. 1881) along with MMPX (Cat. No. 0552) and EHNA hydrochloride (Cat. No. 1261) were purchased from Tocris.

### Microscopy

Methods for immunofluorescent staining have been previously described [14] and adapted for primary mouse cardiomyocytes (ACMs). Briefly, ACMs were plated on laminin coated coverslips (Fisherbrand Microscope Cover Glass, 12-545-80) (1 coverslip per well, 24-well plate) for 1h post isolation. Cells were cultured overnight and treated with drugs as described in the figure legends. Following incubation, cells were washed 1X with PBS (Gibco 14190-144) and fixed for 20min in 4% paraformaldehyde (Electron Microscopy Sciences 15710). Cells were washed 2X with PBS and blocked for 1 hr in 2% Blocker BSA (Thermo 37525). Coverslips were inverted onto 50μL of block solution containing 0.1% Triton X-100 (Sigma T9284) and 1-250 dilution of primary antibodies as indicated on parafilm covered 150mm gridded tissue culture dish (Falcon 353025) and incubated overnight at 4°C. Coverslips were returned to tissue culture dish and washed 3X with block solution and then inverted on 50μL of block solution containing .1% Triton X-100 (Sigma T9284) and 1-250 dilution of secondary antibodies/phalloidin for 1hr at room temperature. Coverslips were then washed 2X with block solution and 1X with PBS and then inverted onto 4μL Vectashield mounting media with DAPI (Vector Laboratories H-1200) and sealed with nail polish (Electron Microscopy Sciences 72180). Cells were imaged on a Zeiss LSM-710 confocal (Carl Zeiss). Micrographs are maximum intensity projections of 12 picture z-stacks average step size of 1.5μm. Projection images were quantitated in Zen Black (Carl Zeiss version 2012 SP5 FP3) from maximum intensity projection by drawing a free polygon outline around the cell and measuring mean fluorescence intensity. Each experiment constitutes at least three biological replicates with at least six individual cells per experiment for a total of at least eighteen total cells quantitated for each data point. The average number of cells per data point is thirty total cells.

### Western blot analysis

Cardiomyocytes were cultured in 12 well plates, with or without insulin, and drugs were added as indicated in the figure legends. Media was then removed, cells were washed with 0.5 mL PBS, and 75 uL of 1X sample buffer containing 25 mM DTT and 1X Halt Protease/Phosphatase Inhibitor Cocktail (ThermoFisher #78442) was added per well. Samples were heated at 95°C for 5 min, resolved by SDS-PAGE (4-12% NuPAGE Bis-Tris gel, Thermo Fisher), transferred to nitrocellulose membranes, and blocked for 30 min with Odyssey TBS blocking buffer (LI-COR). Antibodies were diluted 1:2000 in block buffer and added to blots overnight at 4°C, subsequently washed the following day, and the secondary antibody (IRDye 800CW, LI-COR; 1:10,000 dilution) was incubated for 1h. Following additional washing, blots were imaged on an Odyssey CLx system and analyzed using the Image Studio software (LI-COR).

### Statistical analysis

GraphPad Prism 7.02 was used for statistical analysis and mean fluorescence intensity values were evaluated using one-way ANOVA with multiple comparisons using Tukey’s test. Statistical significance is noted in the figure legends.

## Results

### β-adrenergic signaling is decreased under diabetic conditions in cardiomyocytes

Initial experiments were performed to validate methodology for evaluating PKA signaling by immunofluorescence in adult mouse cardiomyocytes (ACMs). ACMs were isolated from control mice and cultured for 18h in the presence of insulin and then stimulated with isoproterenol (ISO, 0.25μM) for 30min. Cells were then fixed and PKA activity was visualized by immunofluorescence microscopy using an antibody that specifically recognizes the protein consensus phosphorylation sequence (RRXS/T, where S or T is phosphorylated) that is specific for PKA substrates [5]. This antibody is widely used in the literature (132 citations per CiteAb.com) and has been previously used as a means to identify changes in PKA activity by immunohistochemistry [15]. We observed low levels of PKA substrate phosphorylation basally and this was increased approximately 3-fold by ISO treatment (Fig 1A, top panels). Detection of PKA activity by this manner revealed largely diffuse staining but with increased intensity proximal to the sarcolemma and intermittently on Z-bands.

**Fig 1.**
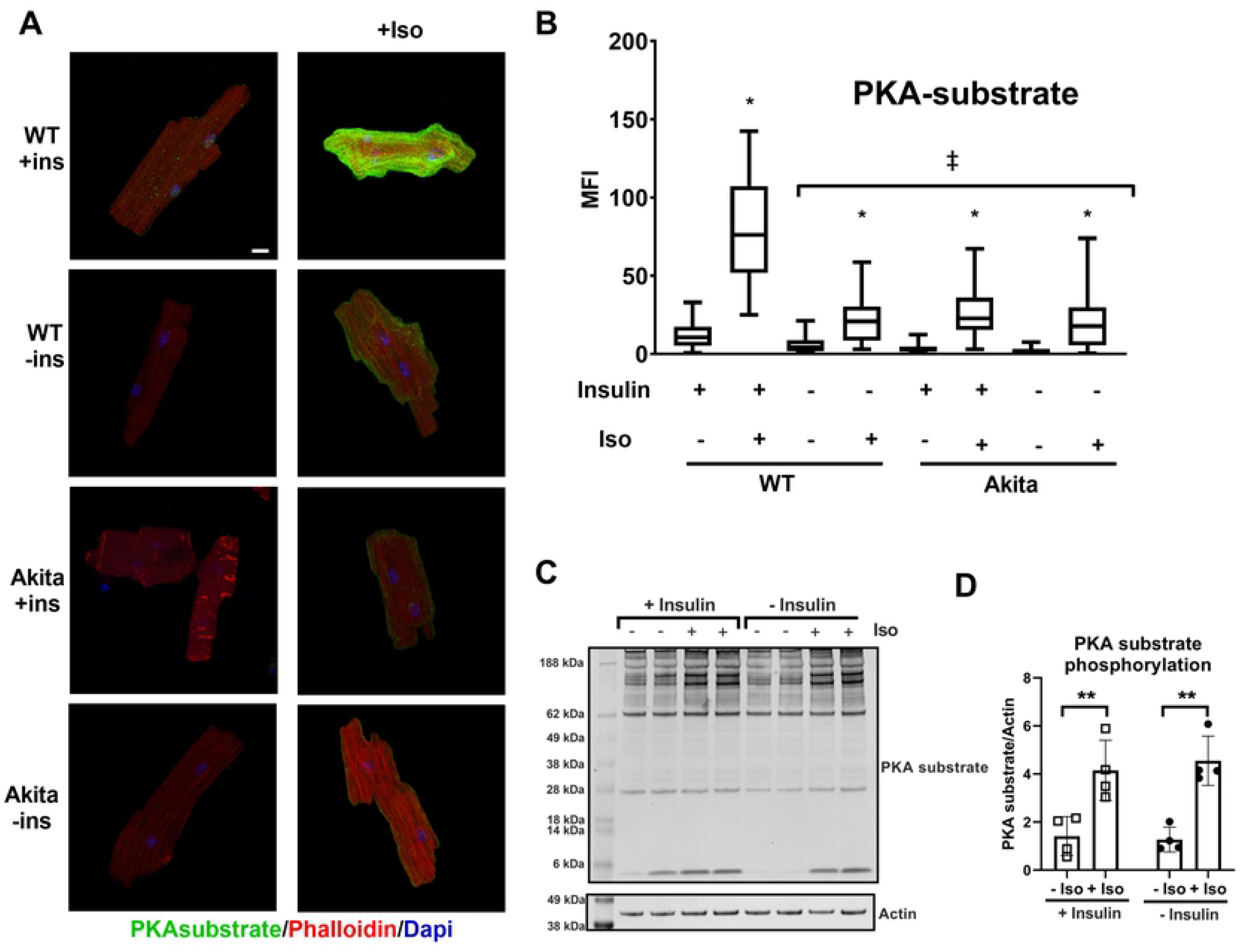
Immunofluorescence detection of PKA signaling reveals a positive role of insulin. (A) Adult mouse cardiomyocytes from wild-type or Akita mice were incubated overnight in the presence or absence of insulin (ins) and treated with 0.25μM isoproterenol for 30min as indicated. Cells were fixed and stained with rabbit anti-PKA substrate antibody visualized with Alexa 488 anti-rabbit secondary and with Alexa 568 labeled phalloidin. Maximum intensity micrographs were acquired as described in Materials and Methods and a representative image for each condition is shown. Scale bar 10μm. (B) Quantitation of mean fluorescence intensity (MFI) for PKA-substrate are presented as whisker plots (n=3 biological replicates, and at least 10 cells per experiment). The box dimensions extend from the 25^th^ to the 75^th^ percentiles; whiskers describe the minimum to maximum values. The median is plotted as a horizontal line within the box. *, Isoproterenol treatment caused a statistically significant increase in all conditions (*p*<.001) by one-way ANOVA. (‡) Isoproterenol stimulated samples from WT without insulin and Akita were significantly reduced (*p*<.001) compared to stimulated WT with insulin by one-way ANOVA with Tukey post hoc test. (C) Representative Western blot for anti-PKA substrate analysis of lysates from primary mouse cardiomyocytes treated as shown. (D) Quantitation of western blot experiments (n=4); **, *p*<.005 by two-way ANOVA with Tukey post hoc test.

We next examined how the lack of insulin affects PKA signaling. Freshly isolated ACMs were cultured in insulin-free media for 18h and then stimulated with ISO. PKA-substrate phosphorylation was significantly blunted both basally and following ISO treatment (Fig 1A). This decrease in PKA-substrate phosphorylation was not due to altered kinetics. A time course study, with increasing durations of ISO stimulation, revealed phosphorylation reached a maximal threshold within 10min regardless of whether insulin was present (S1 Fig). In contrast to the immunofluorescence data, no significant differences were observed when PKA substrate phosphorylation was examined by Western blot (Fig 1B). This suggests unique epitopes are detected by the antibody under conditions that maintain native protein confirmations, as with immunofluorescence detection, as compared to the denaturing conditions of SDS-PAGE.

We next examined whether the chronic hypoinsulemia that occurs with type 1 diabetes is also associated with changes in PKA signaling. Akita mice develop type I diabetes in the absence of obesity and insulitis and is mediated by a mutation in the Ins2 gene [16]. Akita ACMs were isolated and cultured overnight in the presence or absence of insulin. As shown in Fig 1A, Akita ACMs had substantially reduced PKA substrate staining upon stimulation with ISO. Furthermore, culturing cardiomyocytes in the presence of insulin for 18h had no enhancing effect on PKA substrate staining upon ISO stimulation. This supports that decreased insulin signaling affects PKA signaling and that acute insulin treatment of Akita ACMs is insufficient for rescue.

### Insulin signaling is necessary downstream of cAMP production

The decrease in PKA substrate phosphorylation in cardiomyocytes cultured without insulin may be due to changes in β-adrenergic receptors or their response to ligand binding, thereby leading to a dampened response to ISO stimulation. We therefore examined direct activation of PKA using the cell permeable cAMP analog, 8Br-cAMP. Like ISO, 8Br-cAMP induced a robust increase in PKA-substrate phosphorylation in cardiomyocytes isolated from control mice cultured with insulin. However, substrate phosphorylation stimulated by 8Br-cAMP was significantly blunted in ACMs cultured overnight without insulin (Fig 2). Likewise, cardiomyocytes isolated from adult Akita mice had significantly blunted response to 8Br-cAMP and this was not rescued by an 18h culture with insulin. This suggests that the defect in PKA signaling induced by the absence of insulin is downstream of the β-adrenergic receptors and cAMP production.

**Fig 2.**
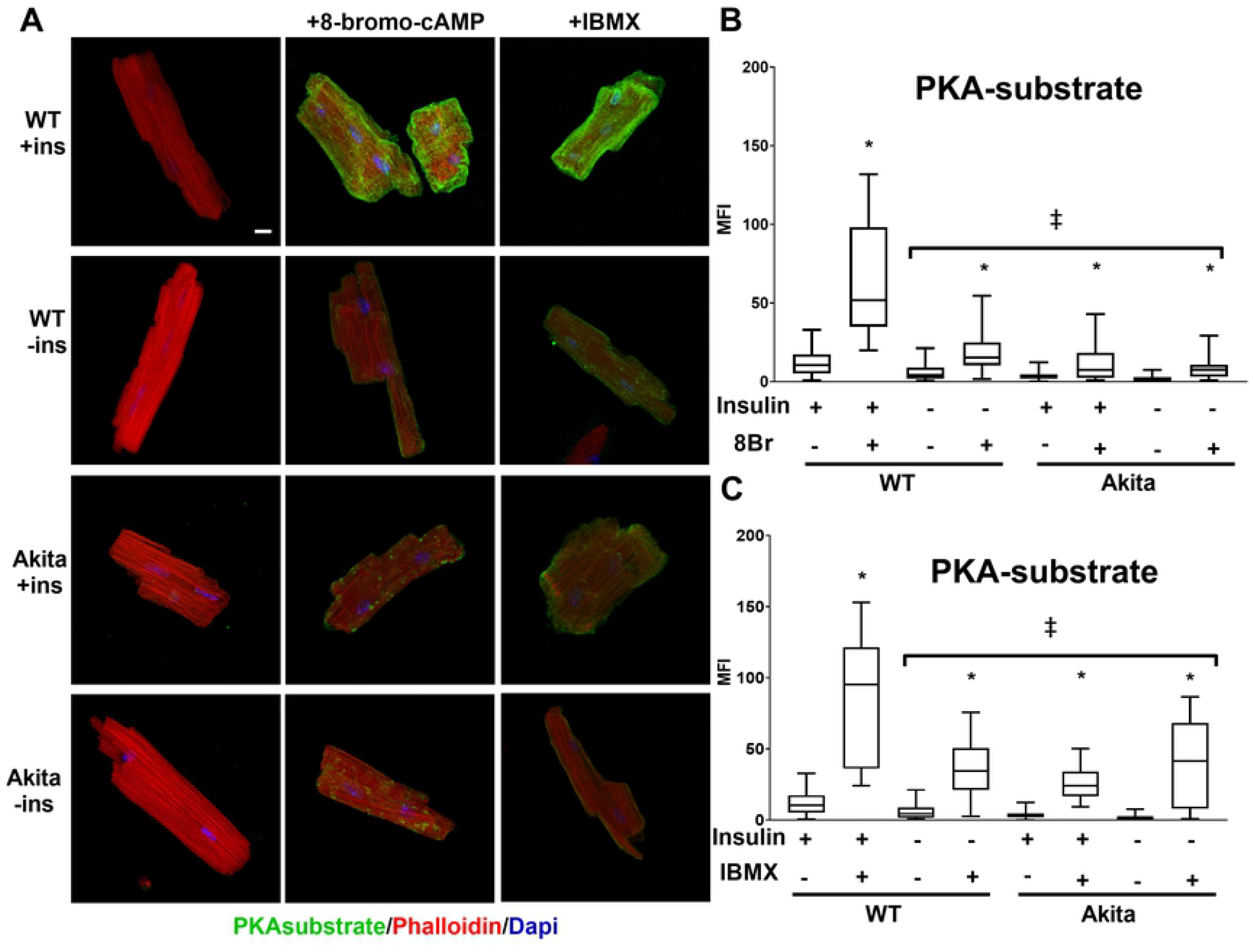
The effect of insulin on PKA signaling occurs downstream of cAMP production. (A) ACMs from wild-type or Akita mice were incubated overnight in the presence or absence of insulin (ins) and treated with 250μM 8-bromo-cAMP (8Br) or 500μM IBMX for 30min as indicated. Cells were fixed and stained with rabbit anti-PKA substrate antibody visualized with Alexa 488 anti-rabbit secondary and with Alexa 568 labeled phalloidin. Maximum intensity micrographs were acquired as described in Materials and Methods and a representative image for each condition is shown. Scale bar 10μm. (B&C) Quantitation of mean fluorescence intensity (MFI) for PKA-substrate are presented as whisker plots, as detailed in Fig 1 (n=3 biological replicates, and at least 6 cells per experiment). (*) Isoproterenol treatment caused a significant increase (*p*<.001) in all conditions. (‡) 8-bromo-cAMP or IBMX stimulated samples from WT without insulin and Akita were significantly reduced (*p*<.001) compared to stimulated WT with insulin. Statistics were performed by one-way ANOVA with Tukey post hoc test.

Alterations in PKA signaling could be attributed to fluctuations in enzyme content. PKA catalytic subunit levels were therefore examined in ACMs from control and Akita mice cultured with or without insulin. As shown in S2 Fig, there were no significant differences in PKA content under the experimental conditions as determined by immunofluorescence or Western blot (not shown). This indicates the loss of PKA substrate phosphorylation is not mediated to decreased PKA catalytic subunit content.

### Phosphodiesterase inhibition increase PKA signaling but do not restore deficits induced by loss of insulin

Phosphodiesterases (PDEs) hydrolyze cAMP and are an essential component in modulating proper PKA signaling. Thus, insulin mediated changes in PDE activity could contribute to the observed effects on PKA signaling shown in Fig 1. We therefore examined the effect of 3-isobuty-1-methylxanthine (IBMX), a nonspecific phosphodiesterase inhibitor, on ACMs from control and Akita mice to determine whether blocking PDE activity is sufficient to recover PKA signaling. Addition of IBMX, in the absence of other PKA agonists, was sufficient to stimulate PKA signaling by 2.5-fold (Fig 2). However, the effect of IBMX was blunted in wild type ACMs cultured without insulin and in Akita ACMs regardless of whether insulin was present. This demonstrates PDE inhibition is sufficient to stimulate PKA substrate phosphorylation similarly to a PKA agonist when insulin is present. However, this effect is blunted by acute insulin withdrawal and in Akita cardiomyocytes.

We next sought to determine the combined effects of ISO and IBMX on PKA signaling. ACMs from control or Akita diabetic mice were cultured overnight in the presence or absence of insulin and then treated with combinations of IBMX and ISO. IBMX enhanced ISO stimulation of PKA substrate phosphorylation under all conditions (Fig 3). This demonstrates the importance of PDE activity in attenuating catecholamine mediated PKA signaling. However, the maximum PKA substrate phosphorylation was nevertheless blunted in control ACMs cultured in the absence of insulin and in Akita cardiomyocytes. Thus, while PDE inhibition enhances PKA substrate phosphorylation under diabetic conditions it does not fully restore it.

**Fig 3.**
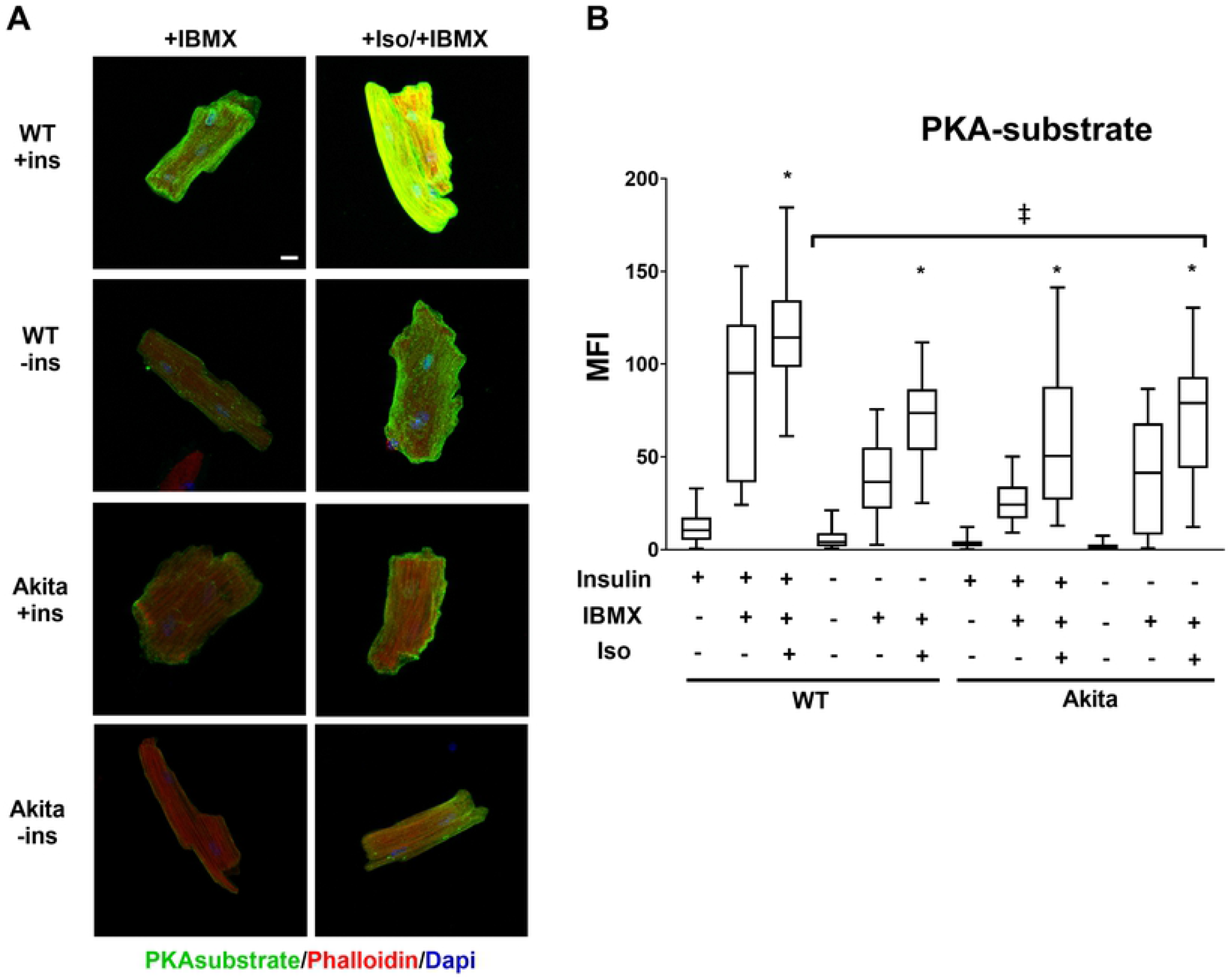
PKA signaling stimulated by the combination of isoproterenol and IBMX is decreased by the absence of insulin. (A) ACMs from wild-type or Akita mice were incubated overnight in the presence or absence of insulin (ins) and treated with 500μM IBMX or .25μM Iso/500μM IBMX for 30min as indicated. Cells were fixed and stained with rabbit anti-PKA substrate antibody visualized with Alexa 488 anti-rabbit secondary and with Alexa 568 labeled phalloidin. Maximum intensity micrographs were acquired as described in Materials and Methods and a representative image for each condition is shown. Scale bar 10μm. (B) Quantitation of mean fluorescence intensity (MFI) for PKA-substrate are presented as whisker plots, as detailed in Fig 1 (n=3 biological replicates, and at least 6 cells per experiment). (*) Isoproterenol and IBMX treatment caused a significant increase (*p*<.001) compared to IBMX alone in all conditions. (‡) stimulated Akita and WT −ins conditions are significantly decreased compared to matching WT +ins conditions. Statistics were performed by one-way ANOVA with Tukey post hoc test.

Recent work has identified a relationship between cardiac insulin signaling and the content and activity of PDE4 [11]. Specifically, the hyperinsulinemia that occurs with type 2 diabetes is associated with increased PDE4B content which thereby decreases cAMP and attenuates β-adrenergic signaling [11]. We therefore tested to see if reciprocally the lack of insulin affects specific phosphodiesterases. Control ACMs were treated with a panel of phosphodiesterase inhibitors including IBMX (nonspecific PDE inhibitor), EHNA (PDE2 specific), MMPX (Calmodulin sensitive cyclic GMP specific), milrinone (PDE3 specific), clostramide (PDE3 specific), RO-20-1724 (PDE4 specific), rolipram (PDE4 specific), and zardavsine (PDE3/4 specific). All of the inhibitors, except EHNA, stimulated phosphorylation of PKA substrates. However, the most pronounced effect was observed with inhibitors of PDE4 (Fig 4). Next, we tested if PDE4 inhibition could restore PKA signaling in the absence of insulin. While PKA signaling in ACMs from control and Akita mice treated with RO-20-1724 closely approximated the effects of IBMX, there was no rescue in ACMs cultured in the absence of insulin (S3 Fig). In addition, we also examined PDE4 content by immunofluorescence and determined that the presence or absence of insulin had no effect on its content (S4 Fig). Consistent with previous reports [17, 18], this supports that PDE4 is the primary regulator of cAMP degradation in mouse cardiomyocytes, that its content is not affected by acute changes in insulin, and that its inhibition is not sufficient to recover PKA activity when insulin signaling is absent.

**Fig 4.**
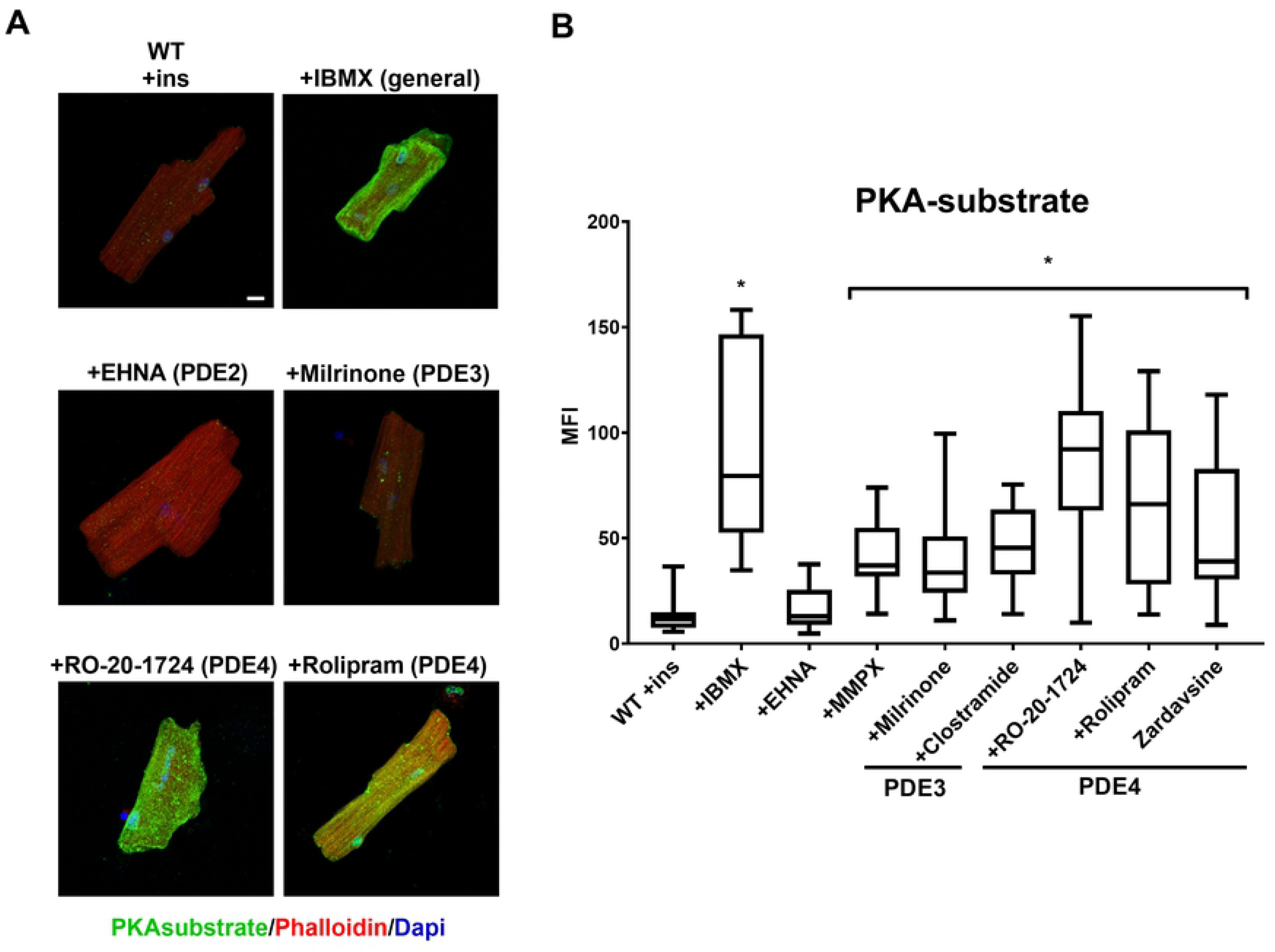
Inhibition of PDE4 increases PKA signaling. (A) ACMs from wild-type or Akita mice were incubated overnight in the presence insulin (ins) and treated with 500μM IBMX or indicated phosphodiesterase inhibitors. Cells were fixed and stained with rabbit anti-PKA substrate antibody visualized with Alexa 488 anti-rabbit secondary and with Alexa 568 labeled phalloidin. Maximum intensity micrographs were acquired as described in Materials and Methods and a representative image for each condition is shown. Scale bar 10μm. (B) Quantitation of mean fluorescence intensity (MFI) for PKA-substrate are presented as whisker plots, as detailed in Fig 1 (n=3 biological replicates, and at least 6 cells per experiment). *, *p*<.001 by one-way ANOVA with Tukey post hoc test.

### Phosphorylation of PFK2 is decreased in diabetic conditions

Our results demonstrate that the absence of insulin affects PKA substrate phosphorylation downstream of β-adrenergic receptors and that this is not mediated by changes in adenylate cyclase or PDE activities. We next evaluated PKA signaling using phospho-specific antibodies to identify substrates that may be differentially phosphorylated in the absence of insulin. The bifunctional enzyme 6-phosphofructo-2-kinase/fructose-2,6-bisphosphatase (PFK-2) is a glycolytic regulator and substrate of PKA [19, 20]. When the cardiac isoform (gene produce of *pfkfb2*) is phosphorylated, PFK-2 increases the production of fructose-2,6-bisphosphate, a potent allosteric activator of the glycolytic enzyme phosphofructokinase-1 (PFK-1). Furthermore, we have previously reported that the content of PFK-2, is regulated by insulin signaling [4, 21]. Control ACMs were cultured for 18h in the presence or absence of insulin and thein stimulated with either ISO or IBMX. As shown in Fig 5, PFK-2 phosphorylation was significantly increased in wildtype ACMs stimulated by either ISO or IBMX regardless of whether cells were cultured with insulin. However, for each condition the magnitude of PFK-2 phosphorylation was significantly less in ACMs cultured in the absence of insulin as compared to those cultured with insulin. In Akita ACMs, PFK-2 phosphorylation was significantly repressed basally and was largely unresponsive to stimulation by either ISO or IBMX. Thus, the immunofluorescence staining of phospho-PFK-2 follows closely to that of PKA substrate antibody.

**Fig 5.**
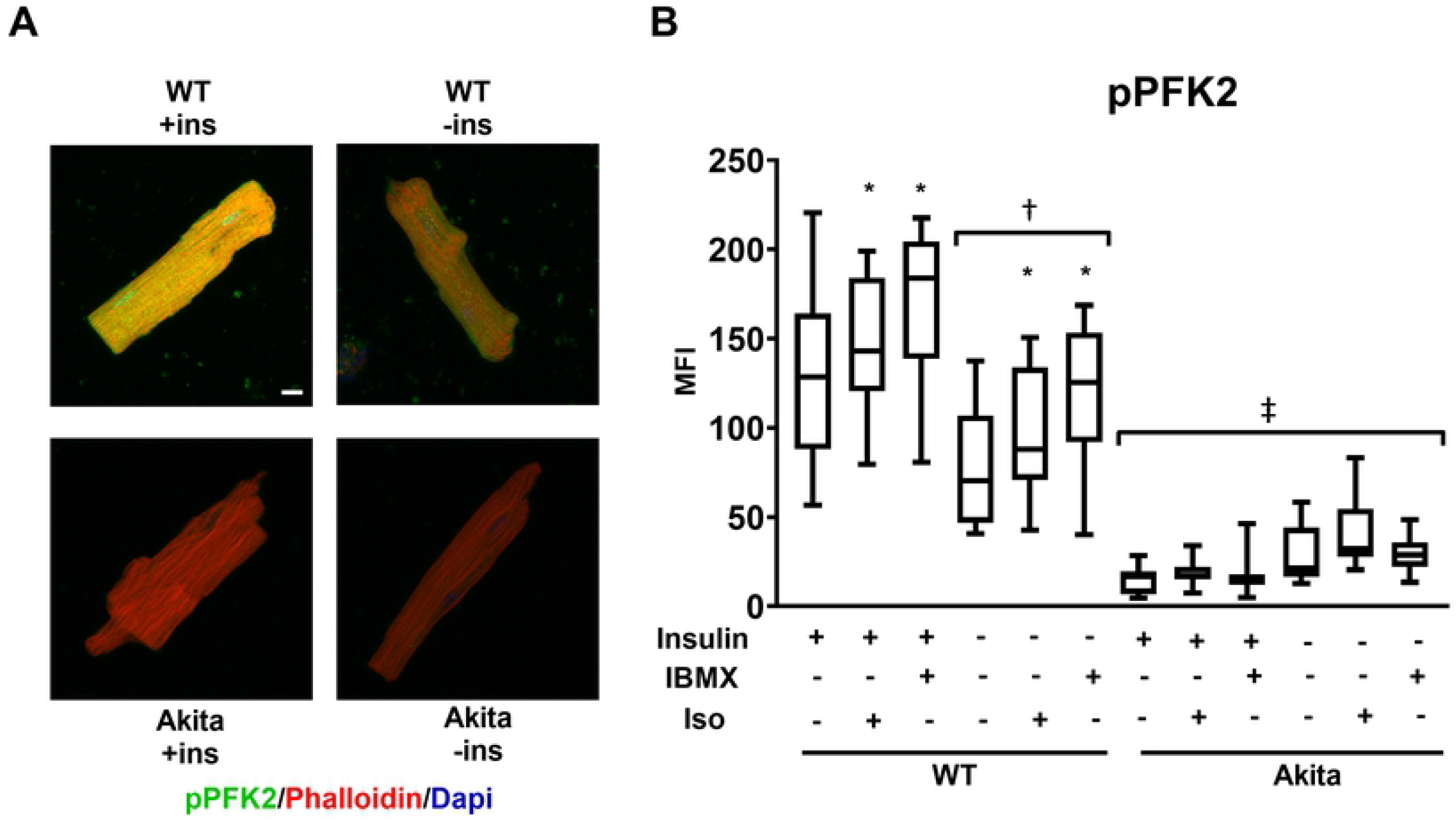
Phosphorylation of PFK2 is decreased in cardiomyocytes cultured in the absence of insulin and with diabetes. (A) ACMs from wild-type or Akita mice were incubated overnight in the presence or absence of insulin (ins) and treated with 500μM IBMX or .25μM Iso for 30min as indicated. Cells were fixed and stained with rabbit anti-phospo-PFK2 antibody visualized with Alexa 488 anti-rabbit secondary and with Alexa 568 labeled phalloidin. Maximum intensity micrographs were acquired as described in Materials and Methods and a representative image for select conditions are shown. Scale bar 10μm. (B) Quantitation of mean fluorescence intensity (MFI) for pPFK2 are presented as whisker plots, as detailed in Fig 1 (n=3 biological replicates, and at least 6 cells per experiment). (*) IBMX or Isoproterenol treatment causes a significant increase (*p*<.001) compared to untreated WT cells; (†) Each WT (−ins) condition is significantly decreased (*p*<.001) compared to WT (+ins); (‡) all Akita conditions are significantly decreased (*p*<.001) compared to comparable WT conditions. Statistics performed by one-way ANOVA with Tukey post hoc test.

We next examined whether the phosphorylation of other well-described PKA substrates exhibit insulin sensitivity. Phospholamban (PBN) is an inhibitor of the sarcoplasmic reticulum calcium dependent ATPase (SERCA2) channel. PBN mediated inhibition of SERCA2 is relieved upon phosphorylation by PKA, thereby increasing calcium cycling in response to β-adrenergic signaling [22]. Troponin I (TnI) is part of the calcium-sensitive troponin complex that decreases myosin-actin crossbridges. Phosphorylation of TnI by PKA decreases the sensitivity of the complex to calcium and is important for increasing the inotropic response [23, 24]. As shown in Fig 6, PBN and TnI were robustly phosphorylated in response to ISO or IBMX and this was unaffected by 18h of insulin starvation. These results demonstrate that there is a differential sensitivity to insulin among β-adrenergic targets, with that metabolic target, PFK-2, being significantly repressed while those involved in contractility sustained.

**Fig 6.**
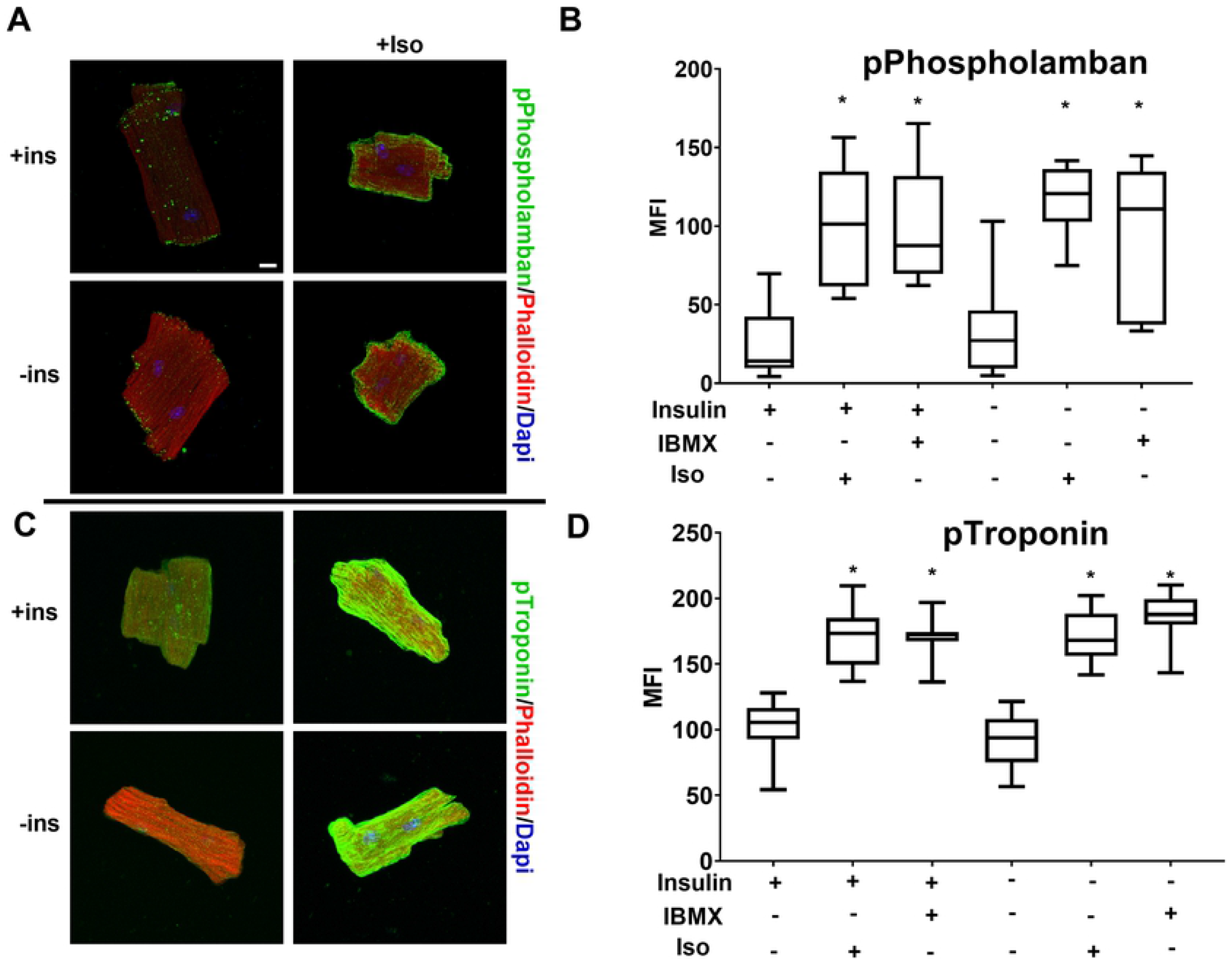
Phosphorylation of β-adrenergic targets Phospholamban and Troponin are unchanged under diabetic conditions. (A&C) Primary mouse cardiomyocytes from wild-type mice were incubated overnight in the presence or absence of insulin (ins) and treated with .25μM Iso or 500μM IBMX for 30min as indicated. Cells were fixed and stained with rabbit anti-phospho-phospholamban (A) or rabbit anti-phospho-Troponin (C) antibody visualized with Alexa 488 anti-rabbit secondary and with Alexa 568 labeled phalloidin. Maximum intensity micrographs were acquired as described in Materials and Methods and a representative image for select conditions are shown. Scale bar 10μm. (B&D) Quantitation of mean fluorescence intensity (MFI) for pPhospholamban (B) or pTroponin (D) are presented as whisker plots, as detailed in Fig 1 (n=3 biological replicates, and at least 6 cells per experiment). (*) IBMX or isoproterenol treatment causes a significant (*p*<.001) increase by one-way ANOVA with Tukey post hoc test.

### PKA substrate phosphorylation remains depressed upon acute insulin administration

Lastly, we sought to determine whether the loss of insulin has a sustained effect on PKA signaling. ACMs from wildtype mice were therefore cultured 18h in the absence of insulin and then stimulated acutely with either ISO, insulin, or in combination. As shown in Figs 7A and 7B, the addition of insulin alone or in combination with ISO failed to rescue PKA signaling under these acute treatment conditions. This effect was not from deficiencies in the insulin signaling pathway. Examination of Akt phosphorylation by Western blot revealed ISO enhanced Akt phosphorylation in ACMs cultured 18h with insulin (Figs 7C and 7D). Acute treatment with additional insulin had minimal effects on Akt phosphorylation, suggesting that desensitization occurs when cells are cultured continuously with insulin. In contrast, ACMs cultured in the absence of insulin showed a robust increase in Akt phosphorylation upon its addition, regardless of the presence of ISO. Thus, the lack of acute effect by insulin on PKA signaling was not due to unresponsiveness of the insulin signaling pathway.

**Fig 7.**
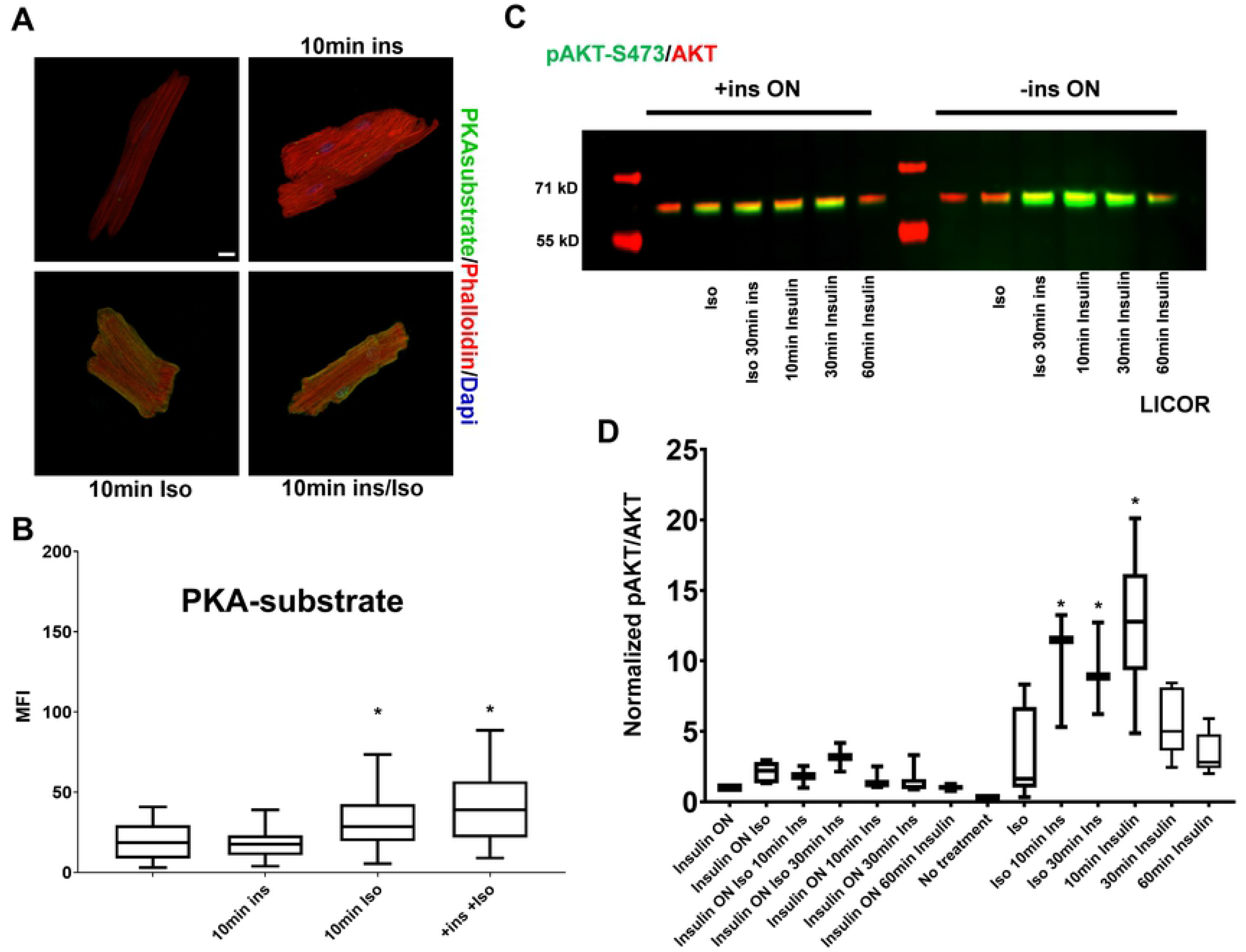
Short term insulin stimulation does not rescue PKA-substrate phosphorylation. (A) Primary mouse cardiomyocytes from wild-type were incubated overnight in the absence of insulin and treated with 0.25μM Isoproterenol or insulin for 10min as indicated. Cells were fixed and stained with rabbit anti-PKA substrate antibody visualized with Alexa 488 anti-rabbit secondary and with Alexa 568 labeled phalloidin. Maximum intensity micrographs were acquired as described in Materials and Methods and a representative image for each condition is shown. Scale bar 10μm. (B) Quantitation of mean fluorescence intensity (MFI) for PKA-substrate are presented as whisker plots, as detailed in Fig 1 (n=3 biological replicates, and at least 6 cells per experiment). (*) Isoproterenol causes a significant increase (*p*<.05) by one way ANOVA. (C) Representative western blot of anti-phospho-Akt-S473 (green) and total Akt (red) analysis of lysates from primary mouse cardiomyocytes treated as in B. (D) Western blot quantitation presented as whisker plots as described in Fig 1. (*) short term insulin stimulation causes a statistically significant increase in AKT phosphorylation at S473 by one way ANOVA with Tukey post hoc test.

## Discussion

β-Adrenergic and insulin signaling pathways are the primary means of modulating moment-to-moment changes in cardiac function and metabolic substrate selection. Nevertheless, interactions between these two pathways are not fully understood. This is important to understand in regard to diabetes where insulin signaling is disrupted and PKA signaling is dysfunctional [3]. In an effort to further understand the interrelationship of these two pathways, we used adult mouse cardiomyocytes as a model system. ACMs have the advantage that they more closely represent in vivo signaling and metabolic conditions as compared to immortalized cardiomyoblasts, such as H9c2 and HL-2 cells [25]. ACMs can also be isolated from different genetic and disease models, as in the Akita mice used here, to interrogate alterations in function at the cellular level. A disadvantage is that ACMs are not amenable to genetic manipulation and have a limited lifespan. For these reasons, we chose to optimize conditions to monitor PKA signaling using an immunofluorescence technique. This approach, to the best of our knowledge, has not been taken before in adult ACMs.

The antibody that we used to measure PKA signaling is validated by the observations that the immunofluorescence intensity is responsive to a β-agonist (ISO), a PKA agonist (8Br-cAMP), and PDE inhibitors. Thus, by this methodology we were able to visualize how insulin affects PKA signaling at the cellular level, specifically in a homogenous cardiomyocyte population, and independently of other systemic factors. Little difference was seen in the subcellular distribution and pattern of PKA substrate phosphorylation in wildtype cells stimulated with either ISO or 8Br-cAMP when cells were culture in the presence of insulin. This suggests that similar pools of PKA were activated by both agonists, resulting in phosphorylation of downstream substrates throughout the cell. When ACMs were cultured in the absence of insulin, though, PKA phosphorylation was substantially blunted in response to all three agonists examined. This was true in both wild type and Akita ACMs. In Akita ACMs, though, the deficiency in PKA signaling was apparent regardless of the presence of insulin. These observations support that the deficiency in PKA signaling was directly at the capacity of PKA to phosphorylate downstream substrates. Interestingly, though, when PKA phosphorylation was examined by Western blot analysis using the same antibody as in the immunofluorescence experiments, no significant differences were seen in the pattern of proteins phosphorylated. The insulin-induced changes in immunofluorescence staining were not due to global changes in ACMs that affected antibody accessibility, though. This is supported by the finding that the phosphorylation of PBN and TnI were robustly induced by agonists even in the absence of insulin. We interpret the results as there being differential recognition of PKA substrate epitopes depending upon the experimental conditions. The immunofluorescence protocol maintains proteins in a native conformation, as compared to Western blot where proteins are denatured.

The effects of insulin on global PKA substrate phosphorylation followed a similar pattern to that observed with phospho-PFK-2 staining. The cardiac isoform of PFK-2 is phosphorylated in response to insulin or β-agonists to increase the levels of fructose-2,6-bisphosphate, an allosteric activator of PFK-1 [26]. This serves to increase glycolytic flux. We have previously reported that PFK-2 content is decreased in the absence of insulin [3]. In the streptozotocin toxic-induced type 1 diabetic model this results in a constitutive decrease in its content. In addition, the PFK-2 that remains is not phosphorylated in response to PKA activation. Our immunofluorescence data follows this similar pattern. In wild type ACMs, PFK-2 is decreased when cultured in the absence of insulin and remains unphosphorylated upon PKA stimulation. In Akita ACMs, a constitutive decrease in PFK-2 is observed. Interestingly, though, culturing Akita ACMs overnight with insulin failed to rescue PFK-2 phosphorylation. This suggests that the mechanisms that normally regulate the expression and activation of PFK-2 are not rescued by a one-time administration of insulin for 18h. Finally, while the pattern of PKA substrate phosphorylation approximates that of phospho-PFK-2, we cannot rule out that other PKA substrates are also affected by insulin.

We demonstrate here that the loss of insulin decreases PKA signaling in ACMs. Reciprocally, previous studies have shown that PKA signaling can also alter the effects of insulin. In neonatal rat cardiomyocytes and mice with chronic β-adrenergic stimulation there is a PKA-dependent decrease in insulin signaling [27, 28]. The mechanism involves insulin receptor desensitization, mediated by Akt [28], and is manifested by a decrease in GLUT4 content and translocation upon insulin stimulation [27]. This PKA-mediated effect contributes to the insulin resistance that is manifested in failing hearts. These role of PKA in modulating insulin signaling, though, are dependent upon the duration of PKA activation. With short-term activation of PKA there is synergistic enhancement of Akt phosphorylation, GLUT4 translocation, and glucose uptake [28, 29]. This mediates the increase in glucose uptake and oxidation in response to acute β-adrenergic stimulation.

Our results provide new insights into how diabetes may impact the heart. We clearly show a decrease in PKA signaling in adult cardiomyocytes when insulin is absent. A novelty of this study was using immunofluorescence microscopy as a means of monitoring PKA signaling. As we show, other methodologies, such as Western blot, would have missed these apparent changes in substrate phosphorylation. Another novel aspect of this study was using primary adult cardiomyocytes isolated from Akita mice. Immortalized cells cannot recapitulate the unique morphology and metabolic aspects of primary cells. Our results support that the deficiency of PKA signaling is not mediated by loss of β-adrenergic receptors, PKA protein, or cAMP production/degradation. Furthermore, the phosphorylation of contractile proteins was similar in control and Akita ACMs, regardless of the presence of insulin. Rather, our results support that a primary defect is the loss of PFK-2 phosphorylation. Future studies must be performed to more exhaustively identify what other substrates may be affected.

## Supporting Information

**S1 Fig. β-adrenergic stimulation increases PKA substrate phosphorylation in adult mouse cardiomyocytes.** (A) ACMs from wild-type mice were incubated overnight in the presence or absence of insulin (ins) and treated with 0.25μM Isoproterenol for 1, 5, and 30min as indicated. Cells were fixed and stained with rabbit anti-PKA substrate antibody visualized with Alexa 488 anti-rabbit secondary and with Alexa 568 labeled phalloidin. Maximum intensity micrographs were acquired as described in Materials and Methods and a representative image for each condition is shown. Scale bar 10μm. (B) Quantitation of mean fluorescence intensity (MFI) for PKA-substrate are presented as whisker plots, as detailed in Fig 1 (n=3 biological replicates, and at least 6 cells per experiment). *, significant difference (*p*<.001) by one-way ANOVA with Tukey post hoc test.

**S2 Fig. PKA catalytic subunit levels are unchanged under diabetic conditions.** (A) ACMs from wild-type or Akita mice were incubated overnight in the presence or absence of insulin (ins) and treated with 0.25μM Isoproterenol or 250μM 8-bromo-cAMP for 30min as indicated. Cells were fixed and stained with rabbit anti-PKA catalytic subunit antibody visualized with Alexa 488 anti-rabbit secondary and with Alexa 568 labeled phalloidin. Maximum intensity micrographs were acquired as described in Materials and Methods and a representative image for each condition is shown. Scale bar 10μm. (B) Quantitation of mean fluorescence intensity (MFI) for PKA-substrate are presented as whisker plots, as detailed in Fig 1 (n=3 biological replicates, and at least 6 cells per experiment).

**S3 Fig. PDE4 inhibition increases PKA signaling.** (A) ACMs from wild-type mice were incubated overnight in the presence or absence of insulin (ins) and treated with 0.25μM Isoproterenol and/or 10μM RO. Cells were fixed and stained with rabbit anti-PKA substrate antibody visualized with Alexa 488 anti-rabbit secondary and with Alexa 568 labeled phalloidin. Maximum intensity micrographs were acquired as described in Materials and Methods and a representative image for each condition is shown. Scale bar 10μm. (B) Quantitation of mean fluorescence intensity (MFI) for PKA-substrate are presented as whisker plots, as detailed in Fig 1 (n=3 biological replicates, and at least 6 cells per experiment). *, significant difference (*p*<.001) by one-way ANOVA with multiple comparisons using Tukey’s test.

**S4 Fig. PDE4D protein levels are unchanged in diabetic or β-adrenergic stimulation conditions.** (A) Primary mouse cardiomyocytes from wild-type (C57-B6) or Akita were incubated overnight in the presence or absence of insulin (ins) and treated with 0.25μM Isoproterenol or 500μM IBMX for 30min. Cells were fixed and stained with rabbit anti-PDE4D antibody visualized with Alexa 488 anti-rabbit secondary and with Alexa 568 labeled phalloidin. Maximum intensity micrographs were acquired as described in Materials and Methods and a representative image for each condition is shown. Scale bar 10μm. (B) Quantitation of mean fluorescence intensity (MFI) for PDE4D are presented as whisker plots, as detailed in Fig 1 (n=3 biological replicates, and at least 6 cells per experiment). *, significant difference (*p*<.001) by one-way ANOVA with multiple comparisons using Tukey’s test.

## References

1. Murtaza G, Virk HUH, Khalid M, Lavie CJ, Ventura H, Mukherjee D, et al. Diabetic cardiomyopathy - A comprehensive updated review. Prog Cardiovasc Dis. 2019;62(4):315–26. Epub 2019/03/30. doi: https://doi.org/10.1016/j.pcad.2019.03.003. PMID: 30922976.

2. Bugger H, Abel ED. Molecular mechanisms of diabetic cardiomyopathy. Diabetologia. 2014;57(4):660–71. Epub 2014/01/31. doi: https://doi.org/10.1007/s00125-014-3171-6. PMID: 24477973.

3. Bockus LB, Humphries KM. cAMP-dependent Protein Kinase (PKA) Signaling Is Impaired in the Diabetic Heart. J Biol Chem. 2015;290(49):29250–8. Epub 2015/10/16. doi: https://doi.org/10.1074/jbc.M115.681767. PMID: 26468277.

4. Bockus LB, Matsuzaki S, Vadvalkar SS, Young ZT, Giorgione JR, Newhardt MF, et al. Cardiac Insulin Signaling Regulates Glycolysis Through Phosphofructokinase 2 Content and Activity. J Am Heart Assoc. 2017;6(12). Epub 2017/12/06. doi: https://doi.org/10.1161/JAHA.117.007159. PMID: 29203581.

5. Shabb JB. Physiological substrates of cAMP-dependent protein kinase. Chem Rev. 2001;101(8):2381–411. Epub 2001/12/26. doi: https://doi.org/10.1021/cr0002361. PMID: 11749379.

6. Depre C, Rider MH, Hue L. Mechanisms of control of heart glycolysis. Eur J Biochem. 1998;258(2):277–90. Epub 1999/01/05. doi: https://doi.org/10.1046/j.1432-1327.1998.2580277.x. PMID: 9874192.

7. Rattigan S, Appleby GJ, Clark MG. Insulin-like action of catecholamines and Ca2+ to stimulate glucose transport and GLUT4 translocation in perfused rat heart. Biochim Biophys Acta. 1991;1094(2):217–23. Epub 1991/09/03. doi: https://doi.org/10.1016/0167-4889(91)90012-m. PMID: 1909899.

8. Brownsey RW, Boone AN, Allard MF. Actions of insulin on the mammalian heart: metabolism, pathology and biochemical mechanisms. Cardiovasc Res. 1997;34(1):3–24. Epub 1997/04/01. doi: https://doi.org/10.1016/s0008-6363(97)00051-5. PMID: 9217868.

9. Riehle C, Abel ED. Insulin Signaling and Heart Failure. Circ Res. 2016;118(7):1151–69. Epub 2016/04/02. doi: https://doi.org/10.1161/CIRCRESAHA.116.306206. PMID: 27034277.

10. Randle PJ, Garland PB, Hales CN, Newsholme EA. The glucose fatty-acid cycle. Its role in insulin sensitivity and the metabolic disturbances of diabetes mellitus. Lancet. 1963;1(7285):785–9. Epub 1963/04/13. doi: https://doi.org/10.1016/s0140-6736(63)91500-9. PMID: 13990765.

11. Wang Q, Liu Y, Fu Q, Xu B, Zhang Y, Kim S, et al. Inhibiting Insulin-Mediated beta2-Adrenergic Receptor Activation Prevents Diabetes-Associated Cardiac Dysfunction. Circulation. 2017;135(1):73–88. Epub 2016/11/07. doi: https://doi.org/10.1161/CIRCULATIONAHA.116.022281. PMID: 27815373.

12. Fu Q, Wang Q, Xiang YK. Insulin and beta Adrenergic Receptor Signaling: Crosstalk in Heart. Trends Endocrinol Metab. 2017;28(6):416–27. Epub 2017/03/04. doi: https://doi.org/10.1016/j.tem.2017.02.002. PMID: 28256297.

13. O’Connell TD, Rodrigo MC, Simpson PC. Isolation and culture of adult mouse cardiac myocytes. Methods Mol Biol. 2007;357:271–96. Epub 2006/12/19. doi: https://doi.org/10.1385/1-59745-214-9:271. PMID: 17172694.

14. Donaldson JG. Immunofluorescence staining. Curr Protoc Cell Biol. 2001; Chapter 4:Unit 4 3. Epub 2008/01/30. doi: https://doi.org/10.1002/0471143030.cb0403s00. PMID: 18228363.

15. Imielski Y, Schwamborn JC, Luningschror P, Heimann P, Holzberg M, Werner H, et al. Regrowing the adult brain: NF-kappaB controls functional circuit formation and tissue homeostasis in the dentate gyrus. PLoS One. 2012;7(2):e30838. Epub 2012/02/09. doi: https://doi.org/10.1371/journal.pone.0030838. PMID: 22312433.

16. Bugger H, Abel ED. Rodent models of diabetic cardiomyopathy. Dis Model Mech. 2009;2(9-10):454–66. Epub 2009/09/04. doi: https://doi.org/10.1242/dmm.001941. PMID: 19726805.

17. Eschenhagen T. PDE4 in the human heart - major player or little helper? Br J Pharmacol. 2013;169(3):524–7. Epub 2013/03/16. doi: https://doi.org/10.1111/bph.12168. PMID: 23489196.

18. Verde I, Vandecasteele G, Lezoualc’h F, Fischmeister R. Characterization of the cyclic nucleotide phosphodiesterase subtypes involved in the regulation of the L-type Ca2+ current in rat ventricular myocytes. Br J Pharmacol. 1999;127(1):65–74. Epub 1999/06/16. doi: https://doi.org/10.1038/sj.bjp.0702506. PMID: 10369457.

19. Kitamura K, Kangawa K, Matsuo H, Uyeda K. Phosphorylation of myocardial fructose-6-phosphate,2-kinase: fructose-2,6-bisphosphatase by cAMP-dependent protein kinase and protein kinase C. Activation by phosphorylation and amino acid sequences of the phosphorylation sites. J Biol Chem. 1988;263(32):16796–801. Epub 1988/11/15. PMID: 2846551.

20. Rider MH, van Damme J, Vertommen D, Michel A, Vandekerckhove J, Hue L. Evidence for new phosphorylation sites for protein kinase C and cyclic AMP-dependent protein kinase in bovine heart 6-phosphofructo-2-kinase/fructose-2,6-bisphosphatase. FEBS Lett. 1992;310(2):139–42. Epub 1992/09/28. doi: https://doi.org/10.1016/0014-5793(92)81315-d. PMID: 1327869.

21. Batushansky A, Matsuzaki S, Newhardt MF, West MS, Griffin TM, Humphries KM. GC-MS metabolic profiling reveals fructose-2,6-bisphosphate regulates branched chain amino acid metabolism in the heart during fasting. Metabolomics. 2019;15(2):18. Epub 2019/03/05. doi: https://doi.org/10.1007/s11306-019-1478-5. PMID: 30830475.

22. MacLennan DH, Kranias EG. Phospholamban: a crucial regulator of cardiac contractility. Nat Rev Mol Cell Biol. 2003;4(7):566–77. Epub 2003/07/03. doi: https://doi.org/10.1038/nrm1151. PMID: 12838339.

23. Zhang R, Zhao J, Mandveno A, Potter JD. Cardiac troponin I phosphorylation increases the rate of cardiac muscle relaxation. Circ Res. 1995;76(6):1028–35. Epub 1995/06/01. doi: https://doi.org/10.1161/01.res.76.6.1028. PMID: 7758157.

24. Zhang R, Zhao J, Potter JD. Phosphorylation of both serine residues in cardiac troponin I is required to decrease the Ca2+ affinity of cardiac troponin C. J Biol Chem. 1995;270(51):30773–80. Epub 1995/12/22. doi: https://doi.org/10.1074/jbc.270.51.30773. PMID: 8530519.

25. Kuznetsov AV, Javadov S, Sickinger S, Frotschnig S, Grimm M. H9c2 and HL-1 cells demonstrate distinct features of energy metabolism, mitochondrial function and sensitivity to hypoxia-reoxygenation. Biochim Biophys Acta. 2015;1853(2):276–84. Epub 2014/12/03. doi: https://doi.org/10.1016/j.bbamcr.2014.11.015. PMID: 25450968.

26. Rider MH, Bertrand L, Vertommen D, Michels PA, Rousseau GG, Hue L. 6-phosphofructo-2-kinase/fructose-2,6-bisphosphatase: head-to-head with a bifunctional enzyme that controls glycolysis. Biochem J. 2004;381(Pt 3):561–79. Epub 2004/06/02. doi: https://doi.org/10.1042/BJ20040752. PMID: 15170386.

27. Mangmool S, Denkaew T, Phosri S, Pinthong D, Parichatikanond W, Shimauchi T, et al. Sustained betaAR Stimulation Mediates Cardiac Insulin Resistance in a PKA-Dependent Manner. Mol Endocrinol. 2016;30(1):118–32. Epub 2015/12/15. doi: https://doi.org/10.1210/me.2015-1201. PMID: 26652903.

28. Morisco C, Condorelli G, Trimarco V, Bellis A, Marrone C, Condorelli G, et al. Akt mediates the cross-talk between beta-adrenergic and insulin receptors in neonatal cardiomyocytes. Circ Res. 2005;96(2):180–8. Epub 2004/12/14. doi: https://doi.org/10.1161/01.RES.0000152968.71868.c3. PMID: 15591229.

29. Stuenaes JT, Bolling A, Ingvaldsen A, Rommundstad C, Sudar E, Lin FC, et al. Beta-adrenoceptor stimulation potentiates insulin-stimulated PKB phosphorylation in rat cardiomyocytes via cAMP and PKA. Br J Pharmacol. 2010;160(1):116–29. Epub 2010/04/24. doi: https://doi.org/10.1111/j.1476-5381.2010.00677.x. PMID: 20412069.

